# Analysis of heat-induced protein aggregation in human mitochondria

**DOI:** 10.1101/253153

**Authors:** Anne Wilkening, Cornelia Rüb, Marc Sylvester, Wolfgang Voos

## Abstract

As proteins in mammalian cells exhibit optimal stability at natural temperatures, small temperature variations may cause unfolding and subsequent non-specific aggregation. As this process leads to a loss of function of the affected polypeptides as well as to further cytotoxic stress, aggregate formation has been recognized as a major pathogenic factor in human diseases. In this study we determined the impact of physiological heat stress on mammalian mitochondria on a proteomic level. The overall solubility of endogenous mitochondrial proteins was only marginally affected by a treatment at elevated temperatures. However, we identified a small subset of polypeptides that exhibited an exceptionally high sensitivity to heat stress. The mitochondrial translation elongation factor Tu (Tufm), a protein essential for organellar protein biosynthesis, was highly aggregation-prone and lost its solubility already under mild heat stress conditions. In parallel, mitochondrial translation as well as the import of cytosolic proteins was defective in heat stressed mitochondria. Both types of nascent polypeptides, derived from translation as well as from import exhibited a strong heat-induced aggregation tendency. We propose a model that a quick and specific inactivation of elongation factors may prevent an accumulation of misfolded nascent polypeptides and thereby attenuate proteotoxicity under stress.

## Introduction

Mitochondria are organelles that fulfill many essential cellular tasks in addition to their traditionally established functions in energy metabolism. These functions are dependent on the enzymatic activities provided by many different mitochondrial polypeptides, of which the majority is encoded in the nucleus while only a few polypeptides are generated by the endogenous mitochondrial protein biosynthesis machinery. The mitochondrial fitness is therefore dependent on a coordinated expression of both nuclear and mitochondrial genes. The mechanisms underlying this coordination process, in particular during adaptation to various stress conditions, are still only inadequately known and are an active field of cell biological research (1).

To maintain mitochondrial functions under normal and stress conditions, a mitochondria-specific protein quality control (PQC) system, consisting of chaperones and proteolytic enzymes, protects the structural integrity of endogenous polypeptides or removes terminally misfolded proteins (2). Apart from a loss of important enzymatic activities, an accumulation of misfolded proteins can also lead to aggregation reactions that result in further structural and functional damage of the cells. As shown in other organelles or in the cytosol, the activity of the PQC system is mainly aimed at the prevention of an accumulation of cellular protein aggregates (3). The proteotoxic effects of aggregates are best exemplified in the occurrence of so-called protein-folding diseases, in particular neurodegenerative diseases that are caused by or at least correlated with the accumulation of specific protein aggregates (4). The chronic expression of aggregation–prone proteins such as pathological poly-glutamine proteins in these diseases declines the protein homeostasis of the cell, which results in further misfolding and aggregation of metastable proteins. Further the overall capacity of protein homeostasis is declined in aging cells, which results in an increased risk of accumulation of toxic protein aggregates (5,6). Research efforts over the past decade have established that mitochondrial dysfunction plays a major role in the etiology of many neurodegenerative diseases (7) as well as in aging processes (8).

Besides mutations and translation errors, there are also environmental stress conditions, which cause protein misfolding and consequently lead to aggregation processes (9). Elevated temperatures or heat stress at non-lethal levels represent a prominent cause of polypeptide denaturation and functional inactivation. As already small increases in temperature can lead to protein unfolding and aggregation, elevated growth temperatures represent a substantial challenge to cellular survival, characterized by a damaged cytoskeleton, fragmentation of organelles, swelling of the nucleus, formation of stress granules and arrest of the cell cycle (10). However, a small increase of the physiological temperature could also be beneficial for organisms under specific circumstances, as fever in vertebrates has a survival benefit by regulating the immune system and protecting the organism against infection (11).

Elevated temperatures activate a cellular heat shock response by inducing the synthesis of heat shock proteins that help to maintain protein homeostasis and ensure the survival of the cell (12). The heat shock response is based on a specific transcriptional response regulated by heat shock transcription factors (HSF) that induce the synthesis of a specific set of molecular chaperones during stress (13). If this protein quality control system is overwhelmed, cells have additional protective mechanisms to deal with aggregated proteins, like sequestration to specific compartments, disaggregation and refolding to a native protein structure, clearance by autophagy as well as asymmetric distribution of aggregates during cell division (9,14).

Although substantial information on the stress-related reactivity of the cytosolic compartment has been obtained, so far there is only little known about the aggregation behavior of endogenous mitochondrial proteins under stress conditions and the effect on the overall mitochondrial function. Previous studies in *S. cerevisiae* have shown that key mitochondrial metabolic enzymes are aggregating and become inactivated under elevated temperatures. The aggregation propensity was modulated by mitochondrial chaperones and the Pim1 protease in the matrix compartment (15). Interestingly, mitochondrial proteins were over-represented in a screen of aging-related protein aggregation in *C. elegans*, connecting mitochondrial protein aggregation with age-related diseases (16).

In this work, we studied the impact of mitochondrial protein aggregation reactions at different stress conditions under *in vivo* like conditions in mitochondria isolated from mammalian cells. We identified aggregation–prone mitochondrial proteins by performing a quantitative proteomic analysis determining the changes in protein solubility after heat stress. In particular, we subjected isolated intact mitochondria to a short heat shock treatment, representing physiological heat stress conditions, to induce protein aggregation and characterize the changes in protein abundance using 2-dimensional differential gel electrophoresis (2D-DIGE). The quantification of the protein spot pattern allowed an identification of temperature-sensitive polypeptides, whose aggregation behavior was further characterized in more detail. Finally, we tested the impact of heat stress on the mitochondrial protein biogenesis reactions by assessing mitochondrial translation as well as the efficiency of polypeptide import.

## Results

### General effects of heat stress on cells and isolated mitochondria

To determine the effects of heat stress on the functional integrity of mitochondria, we measured typical metabolic properties of mitochondria, like the inner membrane potential (ΔΨ), the accumulation of reactive oxygen species (ROS), and the mitochondrial ATP levels. The membrane potential of HeLa cells was analyzed by staining with the potential-dependent mitochondrial dye tetramethylrhodamine (TMRE) and subsequent flow cytometry. As control for a depleted mitochondrial membrane potential, cells were treated with 0.1 μM valinomycin before TMRE staining. We observed that the mitochondrial membrane potential was decreased by about 30% after heat stress at 45°C for 2.5 h (Fig. 1A). The production of ROS was assayed by a staining with the mitochondria-specific superoxide-reactive dye MitoSOXTM. As control we used a treatment with menadione (final concentration (f. c.) 100 μM), a vitamin K3 metabolite that generates superoxide radicals (17). The analysis by flow cytometry showed a decreased production of reactive oxygen species of about 30-40% after incubation at 45°C for 2.5 h compared to unstressed cells (Fig. 1B). We also investigated the temperature dependence of the membrane potential in isolated mitochondria. To monitor the change in membrane potential mitochondria were stained with TMRE and the fluorescence was measured in a microplate reader. Mitochondrial membrane potential was already significantly diminished after 20 min incubation at elevated temperatures. As shown in Fig. 1C membrane potential started to decrease after an incubation at 37°C indicating that isolated mitochondria were more sensitive to increased temperatures as mitochondria in whole cells. At 42°C, representing physiological heat stress, the membrane potential decreased for about 70% and was almost completely abolished after a severe heat treatment at 45°C. We were also interested if there is a change in the ATP levels in isolated mitochondria after incubation at elevated temperatures. Our results showed that there was already a decrease of about 25% in the mitochondrial ATP content after 20 min incubation at 25°C, which was increased after heat shock, resulting in a decrease of more than 80% compared to pre-heat shock levels (Fig. 1D).

**Figure 1.**
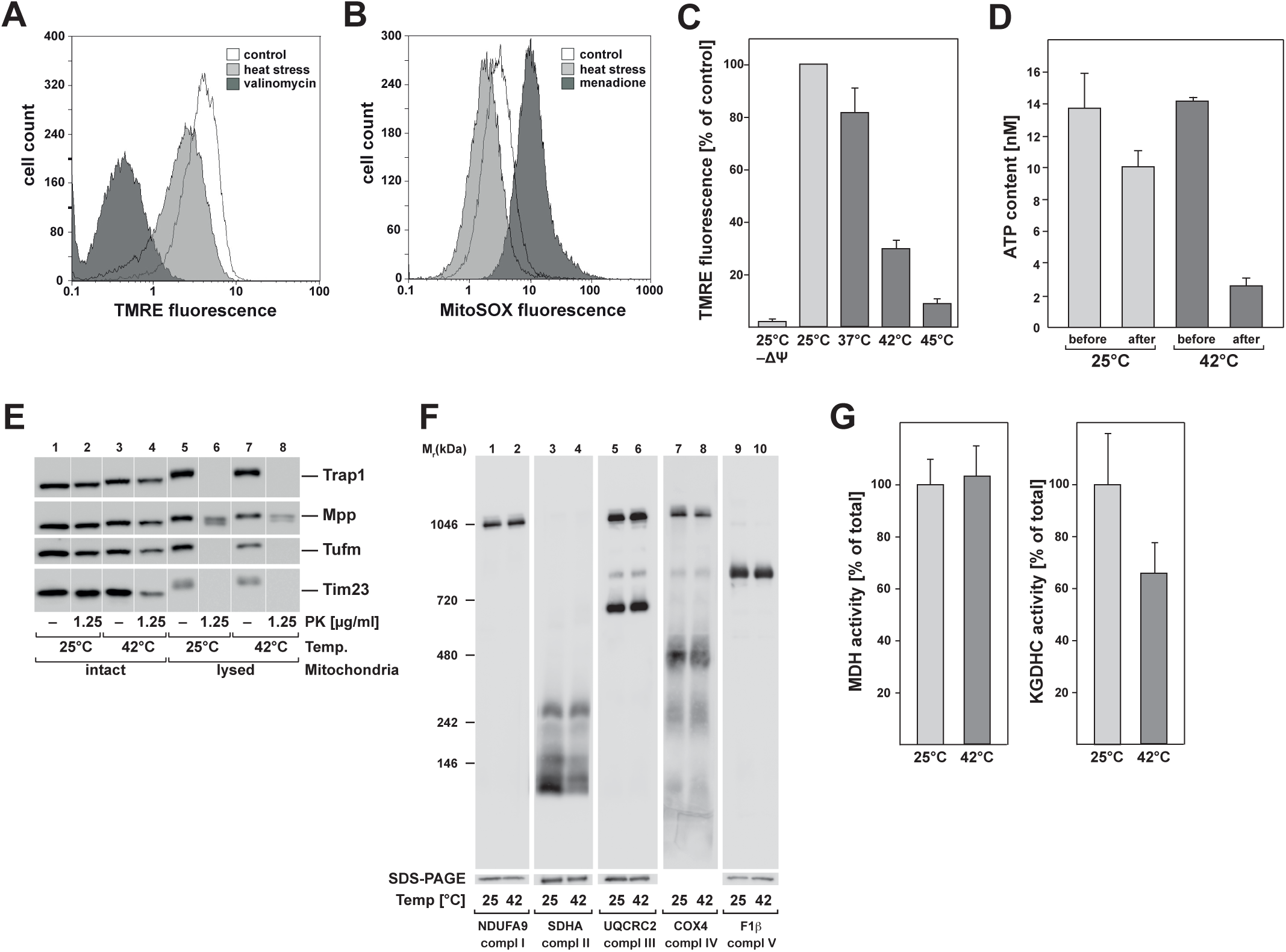
General effects of heat stress on mitochondria. A, B) Flow cytometry analysis of HeLa cells after heat stress. A) Analysis of mitochondrial membrane potential by TMRE staining. Black: negative control (valinomycin treatment), white: control at 37°C, grey: heat stress at 45°C for 2.5 h. B) Detection of reactive oxygen species using the mitochondria-specific superoxide-reactive dye MitoSOXTM. Black: positive control (menadione-treatment), white: control at 37°C, grey: heat stress at 45°C for 2.5 h. Results shown are representatives for two independent experiments. C) Membrane potential of isolated mitochondria after 20 min incubation at different temperatures. The potential was measured by TMRE fluorescence in a microplate reader. Shown are the means of the relative values of the respective sample compared to the control sample at 25°C (n=3) and the standard error of the mean (SEM). D) ATP content in isolated mitochondria after heat shock. The ATP concentration was determined as described in experimental procedures either directly (“before”) or after 20 min incubation at 25°C or 42°C (“after”) E) Integrity of mitochondria after heat stress. Incubation of isolated HeLa wild–type mitochondria at 25°C and 42°C for 20 min and following 20 min treatment with proteinase K (PK) at the indicated concentrations. Mitochondria were lysed prior the treatment with buffer containing 0.5% Triton X-100 as control for a sufficient PK treatment. F) Analysis of the mitochondrial protein complexes in isolated mitochondria after heat stress (20 min, 42°C) compared to unstressed conditions (20 min, 25°C). Mitochondria were analyzed by BN-PAGE or SDS-PAGE and Western blot. Antibodies against subunits of the complex I (NDUFA9), complex II (SDHA), complex III (UQCRC2), complex IV (COX4) and complex V (F1ß) were used. G) Enzymatic activity assays of mitochondrial enzymes after heat shock. Activities of malate dehydrogenase (MDH) and alpha-ketoglutarate dehydrogenase complex (KGDHC) were measured as described after incubation of isolated mitochondria for 20 min at 25°C and 42°C. Values shown are means +/-SEM of the ratio of the activity after heat stress compared to control conditions.

As heat-treated cells exhibited a significant change of the mitochondrial morphology as investigated by fluorescence microscopy (Fig. S1), changing the typical filamentous network structure to a more condensed structure around the nucleus, we investigated also the physical integrity of the heat stressed mitochondria. We used a treatment of isolated intact mitochondria with proteinase K (PK) after a 20 min incubation at 42°C and 25°C (control). PK digests exposed proteins, which are not protected by intact mitochondrial membranes. Western blot analysis showed that there was no digestion of any soluble matrix proteins like Trap1 (heat shock protein 75 kDa), Mpp (mito-chondrial-processing peptidase) and Tufm in both heat stressed and unstressed mitochondria, which suggests that the mitochondrial inner membrane was still intact after heat treatment (Fig. 1E). The translocase of the mitochondrial inner membrane Tim23 was slightly digested by PK in the heat stressed sample indicating that the mitochondrial outer membrane was partially damaged by the incubation at elevated temperatures. To proof that the used conditions were sufficient for protein digestion, we used 0.5% (v/v) Triton X-100 for complete lysis of the mitochondria. All proteins were digested in lysed mitochondria using a PK concentration of 1.25 μg/ml and 20 min incubation, except for Mpp, which was just partially digested.

As one of the main functions of mitochondria is the ATP production by electron transfer through the membrane bound complexes I-V (18), we investigated if there is a structural change in the complexes of the respiratory chain after heat stress. For this purpose the heat shock was performed as described before in isolated mitochondria. Afterwards a separation of the respiratory chain complexes was done under native conditions via Blue-native polyacrylamide gel electrophoresis (BN-PAGE) and analyzed by Western blot using antibodies against subunits of the five different complexes. The experiment showed no significant change in the composition of complexes, despite small changes in complex II and IV (Fig. 1F). We also investigated the enzymatic activities of malate dehydrogenase (MDH) and alpha-ketoglutarate dehydrogenase complex (KGDHC) enzymes as representatives of the tricarboxylic acid cycle (TCA) under heat stress conditions. Activities were tested after 20 min incubation of isolated mitochondria at 42°C compared to 25°C. The MDH activity showed no change after heat shock whereas the KGDHC activity was reduced to 65% of the activity compared with the treatment at 25°C (Fig. 1G), which could indicate a thermo-dependent inactivation of KGDHC.

These results showed that even if the mitochondria and the respiratory chain complexes of the inner membrane are still intact there is a clear temperature-dependent decrease of metabolic functions like an impairment of enzymatic activity and energy production as well as a drop in membrane potential in heat-stressed mitochondria.

### Analysis of the mitochondrial proteome under heat stress

As we have established that heat stress caused functional defects in mitochondria, we determined the effects of heat stress on the structural integrity of the mitochondrial proteome by identifying polypeptides that are prone to aggregate at elevated temperatures. As described above, we treated isolated mitochondria for 20 min at a physiological heat stress temperature of 42°C and 25°C as control. Afterwards mitochondria were lysed by detergent treatment and centrifuged at 125,000 x*g* in order to separate aggregated proteins, which go to the pellet fraction, from soluble proteins, which remain in the supernatant (15,19). To detect quantitative changes in the mitochondrial proteome, we used two-dimensional differential gel electrophoresis (DIGE), which allows the simultaneous separation of three different fluorescently labeled samples on one gel to exclude inter-gel variations. The gels were scanned with dye specific wavelengths and gel matching was performed to detect and quantify the protein spots in the respective samples. Spot detection as well as determination of spot intensities were performed using the Delta2D software (DECODON). For the quantitative analysis, data of two independent experiments, each consisting of triplicates for stressed and unstressed samples were used. On average, a total of 300-400 spots per gel have been detected. The protein spots were manually excised from spots of the Coomassie-stained master gels containing proteins of isolated HeLa mitochondria. Excised spots were digested by trypsin and analyzed by mass spectrometry. In order to obtain a consistent and comparable protein spot pattern, we analyzed the polypeptides present in the supernatant fractions of the different samples. A reduced or absent spot intensity in the supernatant of heat-treated mitochondria in comparison to the control sample would indicate potential aggregation-prone polypeptides that lost their solubility. The short time of incubation at elevated temperature essentially excluded the chance that the reduction of protein amount was caused by degradation processes.

An overlay of the 2D gels obtained after heat-shock treatment allowed a direct comparison of mitochondrial polypeptide solubility. Proteins present in the soluble proteome of the heat-stressed sample were displayed in green, those of the control sample were stained in red resulting in a yellow color overlay picture for protein spots that did not show a change in abundance after the heat shock (Fig. 2A). Therefore red spots would represent proteins present in reduced amounts in the heat stress samples indicating a potential temperature-dependent aggregation. Overall, the abundance of soluble proteins did not change significantly after the heat shock treatment, indicating a high general temperature stability of the mitochondrial proteome, However, there were 11 protein spots, which were consistently reduced in the soluble mitochondrial proteome after heat stress in all six experiments of which 4 proteins could be identified by mass spectrometry (see also Fig. S2). Two of the proteins, Pdia3 (protein disulfide-isomerase A3) as well as Hyou1 (hypoxia up-regulated protein 1), were proteins, which were assigned by the database to be localized in the lumen of the endoplasmatic reticulum (ER). This was likely due to the fact that the used standard method for isolation of mitochondria by differential centrifugation is not able to completely remove all contaminating ER proteins from the mitochondria fraction. However, two of the identified proteins were mitochondrial proteins. The mitochondrial Lon protease homolog (LonP1) showed a partial reduction after heat shock. The most prominent protein spot, which completely disappeared from the supernatant after heat treatment was the mitochondrial elongation factor Tu (Tufm). This result was consistent for all six gels (Fig. 2A, detail sections).

**Figure 2.**
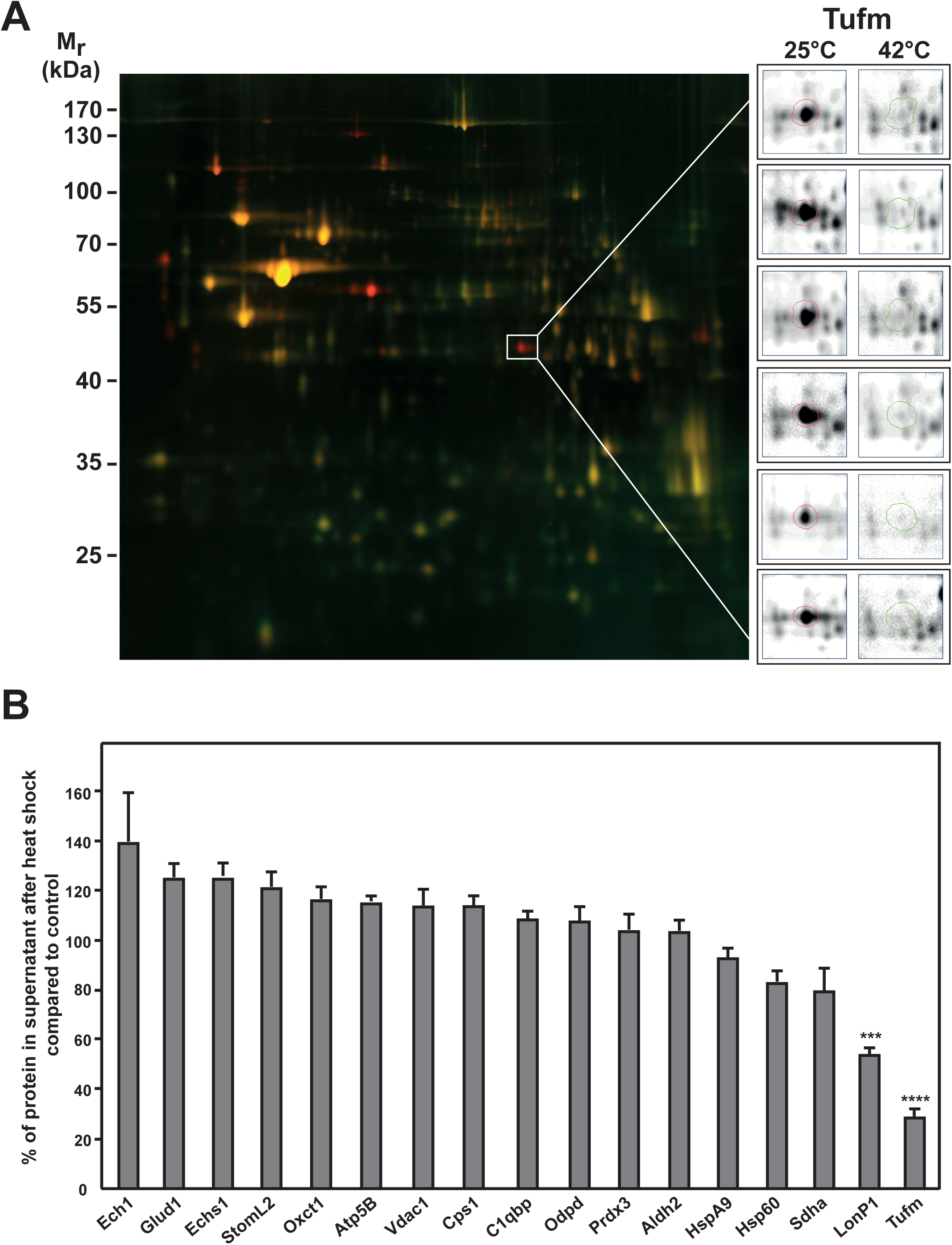
Proteomic analysis of the mitochondrial proteome under heat stress. A) 2D-DIGE spot pattern of heat treated mitochondria. Gel shows an overlay of total soluble protein samples from control (25°C; *red*) and heat-treated (20 min at 42°C; *green*) mitochondria. The gel shown is a representative example of three experiments. Protein spots of the mitochondrial elongation factor Tu (Tufm) are shown enlarged for all six gels analyzed (insets). Mr: relative molecular mass. B) Quantification of selected protein spots. Shown are relative differences of spot intensities between control and heat-stressed mitochondria. Average spot intensities in control samples were set to 100%. Spot detection and quantification were done using the Delta2D software (DECODON). Error bars indicate the standard error of the mean (SEM). The significance was calculated via one-way ANOVA using the Dunnett’s multiple comparisons test. A *p*–value ≤ 0.001 was considered significant and is indicated by asterisks.

We also compared the results for the 17 identified mitochondrial proteins (Table 1; Fig. S2) and quantified the abundance in the soluble proteome after heat shock compared to the control. The spot intensities for each gel were normalized by the total sum of spot intensity at 25°C for each gel (Fig. 2B). There was no significant decrease of abundance in the soluble proteome after heat shock for the majority of proteins, whereas a significant decrease in abundance was observed for the protease LonP1 (about 40%) and Tufm (about 70%) after heat shock. The protein amounts of Sdha (flavoprotein subunit of the succinate dehydrogenase), Hsp60 (60 kDa heat shock protein), and HspA9 (70 kDa heat shock protein) showed only a small decrease of about 5-15%, whereas the other proteins exhibited no significant decrease in the soluble fraction after heat stress compared to the control.

**Table 1.** Quantitative changes in the solubility of mitochondrial proteins under heat stress. Relative amounts of proteins remaining in the supernatant after heat shock were calculated as described in Fig. 2B. Identities of protein spots were analyzed by mass spectrometry in this study and by comparison with previous mass spectrometry results. Protein descriptions were assigned according to database (www.uniprot.org). Molecular weight and pI were calculated using SerialCloner 2-6 by subtracting the transit peptide, which is cleaved after import, from the polypeptide chain, except for Vdac1 as this mitochondrial outer membrane protein exhibits no transit peptide.

| Nr. | Protein | Description | UniProt entry | Size [kDa] | pI | Rel. amount [%] |
| --- | --- | --- | --- | --- | --- | --- |
| 1 | Aldh2 | Aldehyde dehydrogenase | P05091 | 54.34 | 5.76 | 103 ± 4 |
| 2 | Atp5B | ATP synthase subunit beta | P06576 | 51.69 | 4.74 | 115 ± 2 |
| 3 | C1qbp | Complement component 1 Q subcomponent-binding protein | Q07021 | 23.74 | 4.06 | 108 ± 3 |
| 4 | Cps1 | Carbamoyl-phosphate synthetase 1 | A0A024R454 | 160.35 | 6.32 | 113 ± 4 |
| 5 | Sdha | Succinate dehydrogenase [ubiquinone] flavoprotein subunit | P31040 | 67.99 | 6.69 | 79 ± 9 |
| 6 | Ech1 | Delta(3,5)-Delta(2,4)-dienoyl-CoA isomerase | Q13011 | 32.15 | 6.36 | 139 ± 20 |
| 7 | Echs1 | Enoyl-CoA hydratase | P30084 | 28.29 | 5.91 | 125 ± 5 |
| 8 | Glud1 | Glutamate dehydrogenase 1 | P00367 | 55.93 | 7.21 | 125 ± 5 |
| 9 | Hsp60 | 60 kDa heat shock protein | P10809 | 57.88 | 4.96 | 83 ± 4 |
| 10 | HspA9 | Stress-70 protein | P38646 | 68.66 | 5.23 | 93 ± 4 |
| 11 | LonP1 | Lon protease homolog | P36776 | 99.23 | 5.7 | 54 ± 3 |
| 12 | Pdhb | Pyruvate dehydrogenase E1 component subunit beta | P11177 | 35.85 | 5.23 | 108 ± 6 |
| 13 | Prdx3 | Thioredoxin-dependent peroxide reductase | P30048 | 21.5 | 6.14 | 104 ± 6 |
| 14 | Oxct1 | Succinyl-CoA:3-ketoacid coenzyme A transferase 1 | P55809 | 52.01 | 6.33 | 116 ± 5 |
| 15 | StomL2 | Stomatin-like protein 2 | Q9UJZ1 | 35.63 | 5.13 | 121 ± 6 |
| 16 | Tufm | Elongation factor Tu | P49411 | 44.97 | 6.75 | 28 ± 3 |
| 17 | Vdac1 | Voltage-dependent anion-selective channel protein 1 | P21796 | 30.8 | 9.2 | 114 ± 7 |

Overall these results of the proteomic analysis showed that the mitochondrial proteome is relatively resistant against heat stress as the vast majority of mitochondrial proteins are not prone to aggregate. Strikingly one of the mitochondrial proteins, the translation-related protein Tufm, exhibited a very thermolabile behavior by aggregating completely under heat stress.

### Aggregation behavior of mitochondrial elongation factors during stress

Having determined that Tufm is the most prominent protein disappearing from the soluble fraction under heat stress conditions, we wanted to confirm if this behavior was due to protein aggregation and which relevance a potential aggregation has on the functionality of affected mitochondria. In order to get more information about a temperature-dependent aggregation of Tufm in comparison to other mitochondrial proteins, we incubated isolated mitochondria at control temperature (25°C) as well as under mild and strong heat shock conditions (37°C, 42°C, and 45°C). Although a temperature of 37°C can be considered as physiological in HeLa cells, our results (Fig. 1C) have indicated that isolated mitochondria showed already some temperature-sensitivity under this condition.

Isolated HeLa mitochondria were lysed after heat treatment and aggregated proteins (pellet) were separated from the soluble proteins (supernatant) by high-speed centrifugation and analyzed by SDS-PAGE and Western blot. In addition to Tufm, the aggregation behavior of the elongation factor Ts (Tsfm), which is associated with Tufm during translation inducing GDP to GTP exchange (20), as well as the mitochondrial 39S ribosomal protein 38 (Mrpl38) was investigated. A significant amount of Tufm and Tsfm as well as Mrpl38 was recovered in the pellet already at 37°C and their aggregation tendency increased further with higher temperatures (Fig. 3A). Notably, at 42°C Tufm and Mrpl38 were not detected any more in the supernatant fraction, indicating a complete aggregation. The mitochondrial matrix proteins Aco2 (aconitate hydratase) and Hsp60 also started to aggregate at 37°C, but not as strong as Tufm. Whereas Aco2 was completely aggregated after incubation at 45°C, part of Hsp60 stayed soluble. The translocase of the inner mitochondrial membrane Tim23 served as a control, which showed no temperature-dependent aggregation.

**Figure 3.**
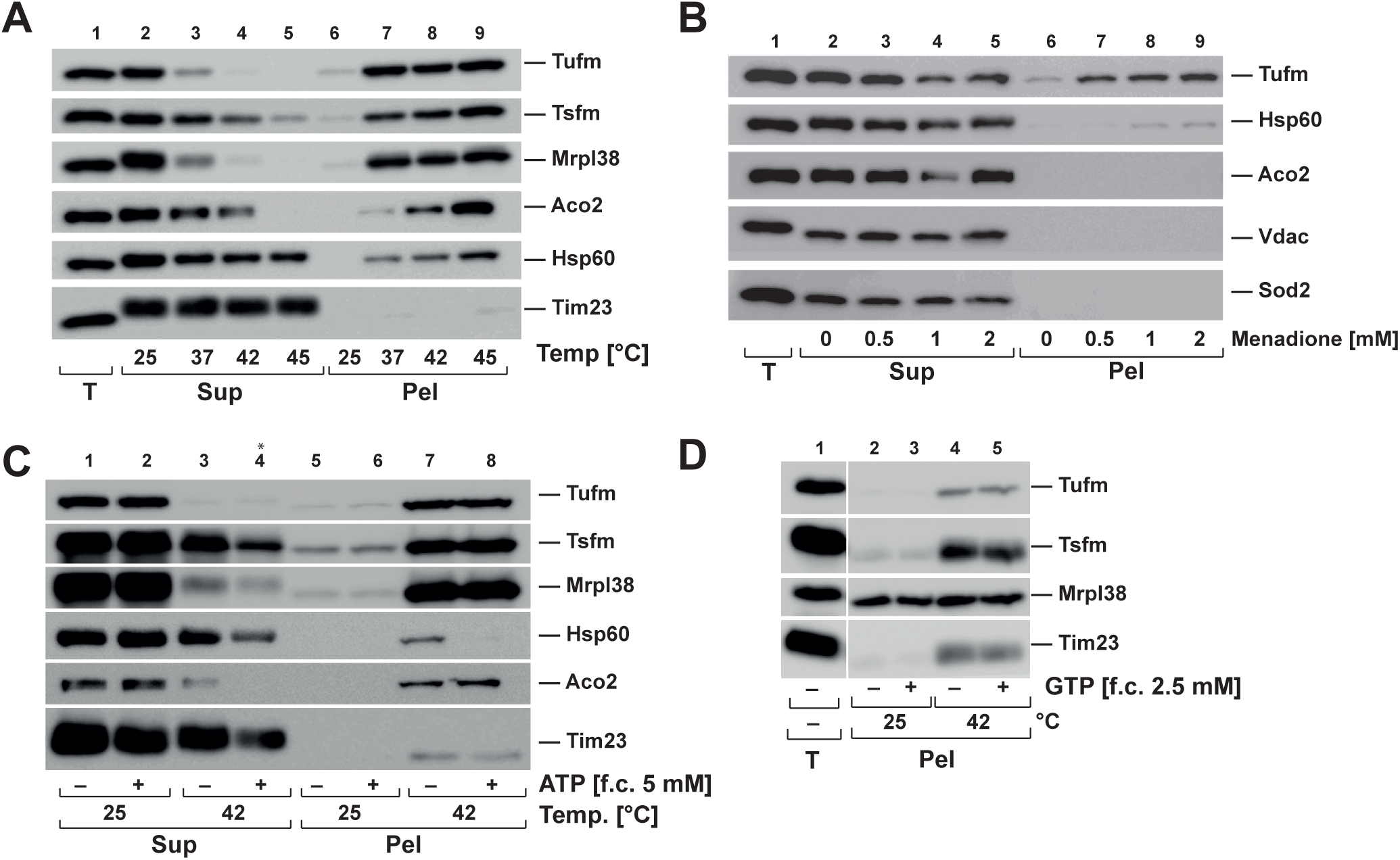
Aggregation behavior of mitochondrial elongation factors during stress. A) Aggregation under heat stress. Isolated mitochondria were incubated for 20 min at indicated temperatures and aggregates were separated by centrifugation at 20,000 x*g*. Total lysates (T), supernatants (Sup) and pellets (Pel) were analyzed by SDS-PAGE and Western blot. B) Aggregation under oxidative stress. Isolated mitochondria were treated with menadione as indicated for 20 min and analyzed as described above. C) ATP dependence of Tufm aggregation. Isolated mitochondria were heat stressed for 20 min at 42°C in presence and absence of ATP (f. c. 5 mM) and analyzed as described above. Asterisk in lane 4 indicate that less sample was loaded due to loss of sample material during the procedure. D) GTP dependence of Tufm aggregation. Isolated mitochondria were heat stressed in presence and absence of GTP (f. c. 2.5 mM) and analyzed as describe above.

To investigate if other stress conditions such as the formation of reactive oxygen species also induced an aggregation of mitochondrial proteins, mitochondria were treated with menadione. Isolated mitochondria were incubated with different menadione concentrations and aggregated and soluble proteins were separated and analyzed as described above. Under oxidative stress conditions we observed no aggregation for the mitochondrial matrix proteins Aco2 and Sod2 (superoxide dismutase) as well as for the outer membrane control protein Vdac (voltage-dependent anion-selective channel protein) (Fig. 3B). Whereas just a small subset of Hsp60 aggregated after treatment with high menadione concentrations, significant amount of the elongation factor Tufm aggregated in a concentration-dependent manner, starting at 0.5 mM menadione. However, compared to the temperature behavior of Tufm, there was still a substantial amount of soluble protein detected in the supernatant, indicating that Tufm may be more sensitive to heat than to oxidative stress.

Molecular chaperones are the most prominent cellular enzymes involved in aggregation prevention and refolding of heat-denatured proteins (21). As most molecular chaperones require ATP for their activity, we wanted to determine if there is an ATP-dependence of Tufm aggregation. Isolated HeLa mitochondria were heat stressed at 42°C in the presence and absence of ATP. Aggregates and soluble proteins were isolated and analyzed as described before. It was shown that although there was an ATP-effect after heat shock visible for the shock protein Hsp60, which was more stable in the presence of ATP, there was no change of the Tufm as well as Tsfm, Mrpl38 and Aco2 aggregation (Fig. 3C), indicating that at least under the chosen conditions there is no aggregation-protective effect for the analyzed proteins mediated by mitochondrial chaperones.

Tufm is a GTPase, which hydrolyzes GTP as energy source for conformational changes in the process of transferring the aminoacyl-tRNA to the A-site of the ribosome (20). As the presence or absence of guanine nucleotides might influence protein stability of Tufm, we also determined the influence of GTP on its temperature-dependent aggregation. Isolated mitochondria were heat stressed in absence and presence of GTP and aggregates were isolated and analyzed as described above. As shown in Fig. 3D there was no GTP dependence of the aggregation of Tufm at 42°C as well as for Tsfm. The ribosomal protein Mrpl38 was already present in the pellet fraction at 25°C in the same amount as after heat shock and there was no visible change in the presence of GTP.

### Time-dependence of aggregation under heat stress

The kinetics of the mitochondrial protein aggregation was studied by incubation of isolated mitochondria for several time points from 0 to 60 minutes at heat stress and control conditions. Aggregated proteins were analyzed by SDS-PAGE and Western blot. The percentage of aggregation was calculated by the amount of aggregated protein compared to the total protein amount (Fig. 4). The mitochondrial elongation factors Tufm and Tsfm both exhibited remarkable fast aggregation kinetics. These proteins started to aggregate already after 2 min incubation at 42°C and practically reached a maximal level already after 5 min. The protein amounts in the pellet did not increase significantly over time any more, indicating a very thermolabile behavior. The ribosomal protein Mrpl38 also showed fast aggregation kinetics with around 50% aggregation after 5 min, which increased over time up to 70%. However 10% of Mrpl38 was already detected in the pellet fraction without any heat stress indicating a temperature independent tendency to sediment under the used centrifugation conditions, most likely due to its association with ribosomal particles. It should be noted that Tufm, Tsfm, and Mrpl38 showed also some aggregation at control conditions (25°C) after longer incubation times indicating an intrinsic low stability. In contrast to the former proteins, Aco2 and Hsp60 showed slower aggregation kinetics after heat shock. Whereas just 15-20% of these proteins were found aggregated after 5 min, the amount of aggregated protein increased over time up to 90% (Aco2) and 60% (Hsp60) after 60 min heat shock. Tim23 served as a control for a heat-stable protein that only showed a minor aggregation propensity (less than 20%) even after 1 h of heat shock. These observations indicate that in particular the translation factors Tufm and Tsfm are extremely thermo-sensitive and aggregate very quickly under heat shock conditions.

**Figure 4.**
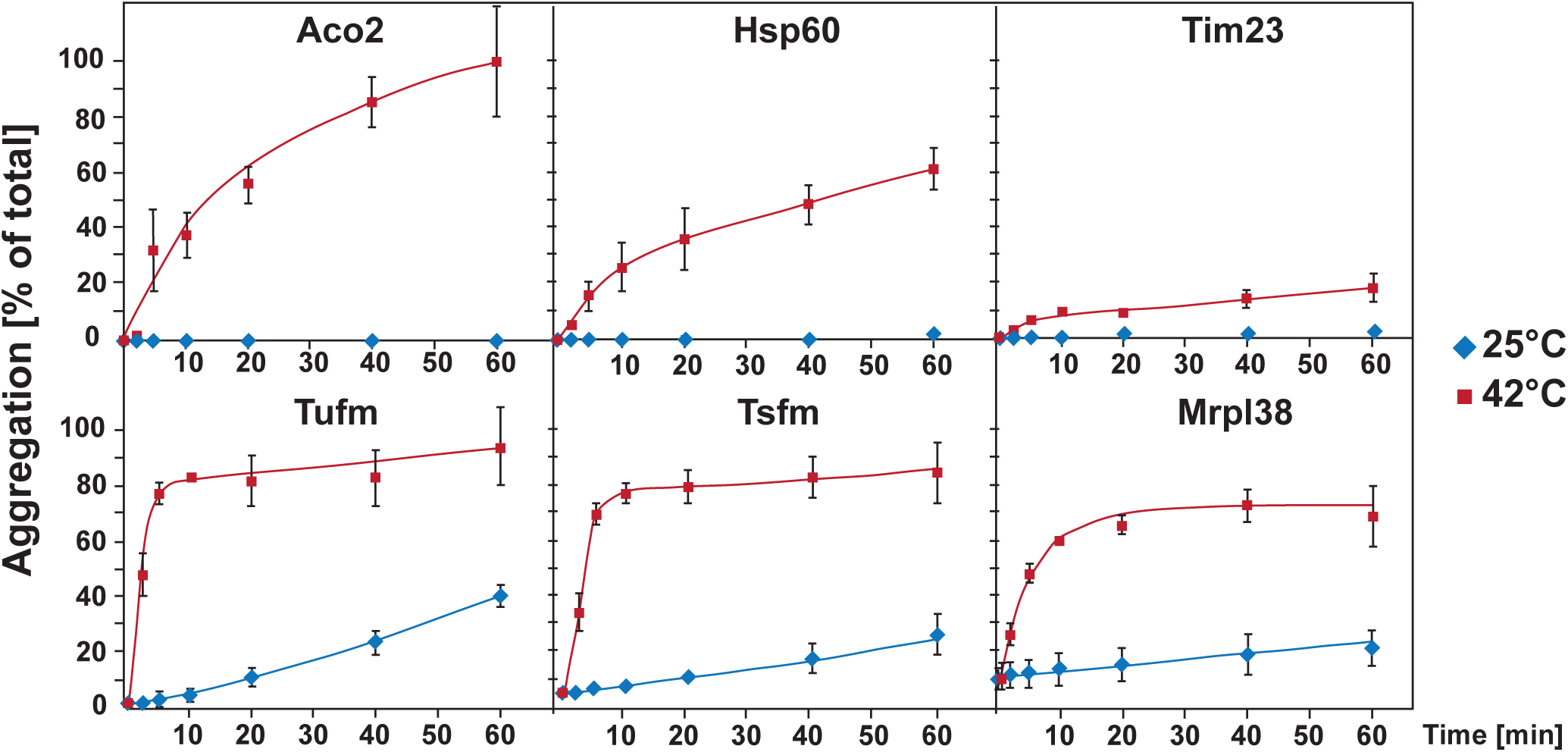
Time dependence of aggregation under heat stress. Time dependence of aggregation of mitochondrial proteins after heat stress. After incubation of isolated mitochondria at 42°C for indicated times aggregates were analyzed by SDS-PAGE and Western blot. Percentage of aggregation was calculated by the ratio of aggregated protein (pellet) compared to the total protein (supernatant + pellet). Shown are mean values of three experiments; error bars indicate the standard error of the mean (SEM).

### Properties of Tufm aggregates

In order to analyze the nature of the Tufm aggregates we analyzed their sedimentation behavior by differential centrifugation. Isolated mitochondria were subjected to a heat treatment as described before and after lysis of the mitochondria aggregated proteins were fractionated at different centrifugal forces to get information about size and density of the formed aggregates. Whereas Tufm and Tsfm were soluble at control conditions and just small amounts were detected in the pellet fraction after 125,000 x*g*, after heat shock aggregates were already detected after a low spin step (5 min at 1600 x*g*) suggesting that they represent rather big aggregates (Fig. 5A, left panel). After 20 min centrifugation at 20,000 x*g* the Tufm aggregates were completely sedimented, which caused us to keep this as standard condition for isolating aggregates in the following experiments. As it could be assumed that elongation factors might also sediment due to a possible attachment to the ribosome in the process of translation, we compared the sedimentation behavior of Tufm with the ribosomal protein Mrpl38. In contrast to Tufm, at control conditions (25 °C) Mrpl38 was mainly detected in the pellet fraction after 40 min centrifugation at 125,000 x*g*. This indicated that the presence of Mrpl38 in the pellet is due to the sedimentation of whole ribosomal subunits after high-speed sedimentation. However, after heat stress also Mrpl38 partially aggregated, as indicated by sedimentation already at lower centrifugation speeds. Still a significant amount of Mrpl38 could be found in the putative ribosome pellet, while none of the elongation factor components were present in this fraction, at least under heat stress conditions. Hence, the sedimentation behavior of Tufm and Tsfm represented a genuine aggregation phenomenon independent of the behavior of ribosomal proteins. This result was confirmed by analyzing the heat-stress aggregates by sucrose gradients, which also showed an independent aggregation behavior of Tufm and Mrpl38 (Fig. S3) as well as there was no influence of high salt concentrations on the heat-induced Tufm aggregation detected indicating no ionic binding between the protein and the ribosome present (Fig. S4).

**Figure 5.**
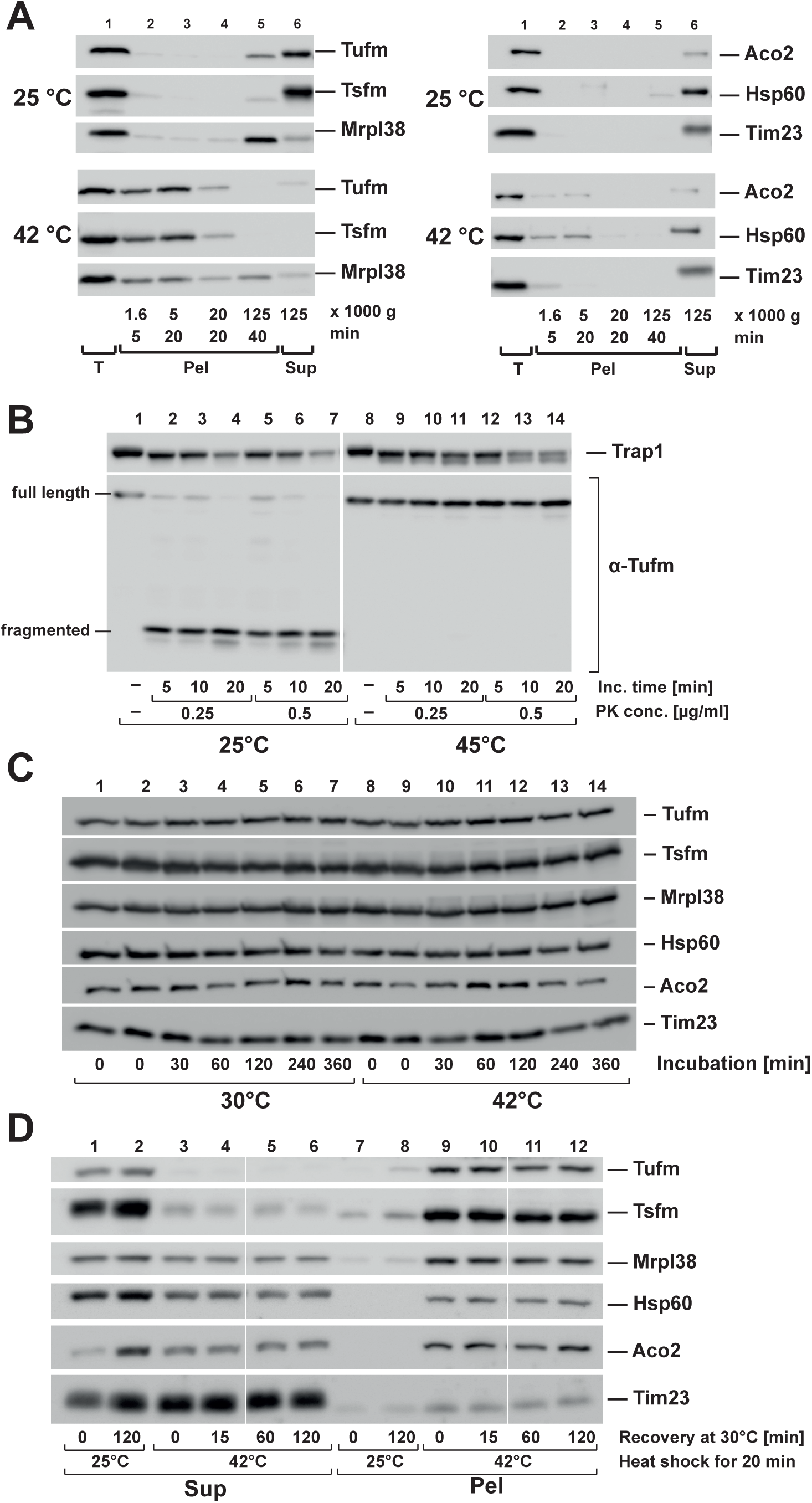
Nature of Tufm aggregates after heat stress. A) Analysis of aggregate size after heat stress. Isolated mitochondria were incubated for 20 min under stress (42°C) and control (25°C) conditions. Protein aggregates were fractionated by differential centrifugation (supernatants of each centrifugation step were transferred to the next centrifugation step). B) Protease resistance of Tufm aggregates. Isolated mitochondria were incubated for 20 min at the indicated temperatures. Mitochondria were lysed with 0.5% Triton X-100 and the extracts treated with the indicated concentrations of proteinase K. The resulting proteolytic fragments of Tufm as well as a matrix control protein (Trap1) were detected by SDS-PAGE and Western blot. C) Degradation of mitochondrial proteins during heat stress. Intact energized mitochondria were incubated for up to 6 h at 30°C (control) and 42°C (heat stress). Protein was precipitated with TCA and analyzed by SDS-PAGE and Western blot using the indicated antibodies. D) Recovery of aggregation after heat shock. Isolated mitochondria were heat stressed at 42°C for 20 min and afterwards incubated for up to 2 h at 30°C. During the recovery incubation, mitochondria were energized and supplied with ATP. Supernatants (Sup) and pellets (Pel) were separated by centrifugation and analyzed by SDS-PAGE and Western blot.

Although the less thermo-sensitive proteins Aco2 and Hsp60 were still partial soluble after heat stress, the stress-induced aggregates were also found already in the low speed pellets (5 min centrifugation at 20,000 x*g*). The control protein Tim23 was not aggregating after heat stress (Fig. 5A, right panel). That the pelleted material of Tufm represented genuine aggregates was confirmed by their resistance against the proteolytic digestion of the proteinase K. Heat-treated mitochondria were lysed with the non-denaturing detergent Triton X-100 and then incubated with different proteinase K concentrations. Whereas Tufm under control conditions (25 °C) was completely digested to a smaller fragment already at very low protease concentrations, it remained completely resistant against digestion after heat shock (Fig. 5B). In comparison, the mitochondrial matrix protein Trap1 was only partially digested by proteinase K both under control and stress condition, indicating an intrinsic protease resistance. An investigation of the mitochondrial protein levels over 6 h under heat stress incubation (42°C) and control (30°C) conditions showed that there was no change in the total protein amounts (Fig. 5C). This verified that the decrease in the soluble amounts of the proteins Tufm and Tsfm was indeed caused by a heat–induced aggregation and not due to proteolytic degradation.

As the elongation factor Tufm exhibited a strong aggregation propensity after heat stress, we wanted to investigate if the solubility of the protein could be recovered over time after the heat stress has ceased. For this purpose heat-stressed mitochondria were subjected to a recovery incubation at 30°C for 15, 60 and 120 min. During recovery mitochondria were supplied with an ATP regenerating system and mitochondrial substrate molecules to keep them fully energized. Aggregates and soluble proteins were separated as described before by centrifugation and analyzed by SDS-PAGE and Western blot. In the tested time range no re-solubilization of aggregated Tufm nor of the other tested, less aggregation-prone, proteins Aco2 and Hsp60 was observed (Fig. 5D), indicating that the aggregated polypeptides could not be quickly disaggregated.

### Mitochondrial protein biosynthesis under heat stress

As the elongation factor Tufm promotes the binding of aminoacyl-tRNA to the A-site of the ribosome during protein translation (22), we wanted to investigate the correlation between the thermal aggregation of Tufm and its effect on mitochondrial translation reactions. We investigated the impact of heat stress on the translation efficiency in isolated mitochondria by checking radiolabeled translation products after different times of pre-incubation of isolated mitochondria at 42°C. Newly synthesized proteins encoded in the mitochondrial genome were detected by an incubation with ^35^S-labeled amino acids methionine/cysteine for 45 min (Fig. 6A right panel). The labeled protein bands could be assigned to subunits of mitochondrial respiratory chain complexes NADH dehydrogenase, Cytochrome *c* oxidase, Cytochrome *bc*1 complex and ATP-Synthase (23) (Fig. 6A left panel). Control samples were treated with cycloheximide or chloramphenicol to inhibit translation reactions. After two minutes of heat stress the amount of newly translated proteins is comparable to the control conditions whereas with increasing incubation times at 42°C the amount of translation products decreased significantly (Fig. 6A). As shown for control and heat stressed samples both cycloheximide and chloramphenicol blocked the mitochondrial translation completely (Fig. 6A). A quantification of the overall translation efficiency (sum of total radioactivity per lane) revealed that there is a clear correlation between the time of heat stress and the decline of mitochondrial translation efficiency (Fig. 6B). While after 2 min of heat stress the level of freshly translated proteins was still comparable to the control conditions, translation efficiency decreased to less than 20% after 20 min. It was also shown that the *in organello* translation after 20 min heat stress could not be recovered after two hours at 25°C (Fig. S5).

**Figure 6.**
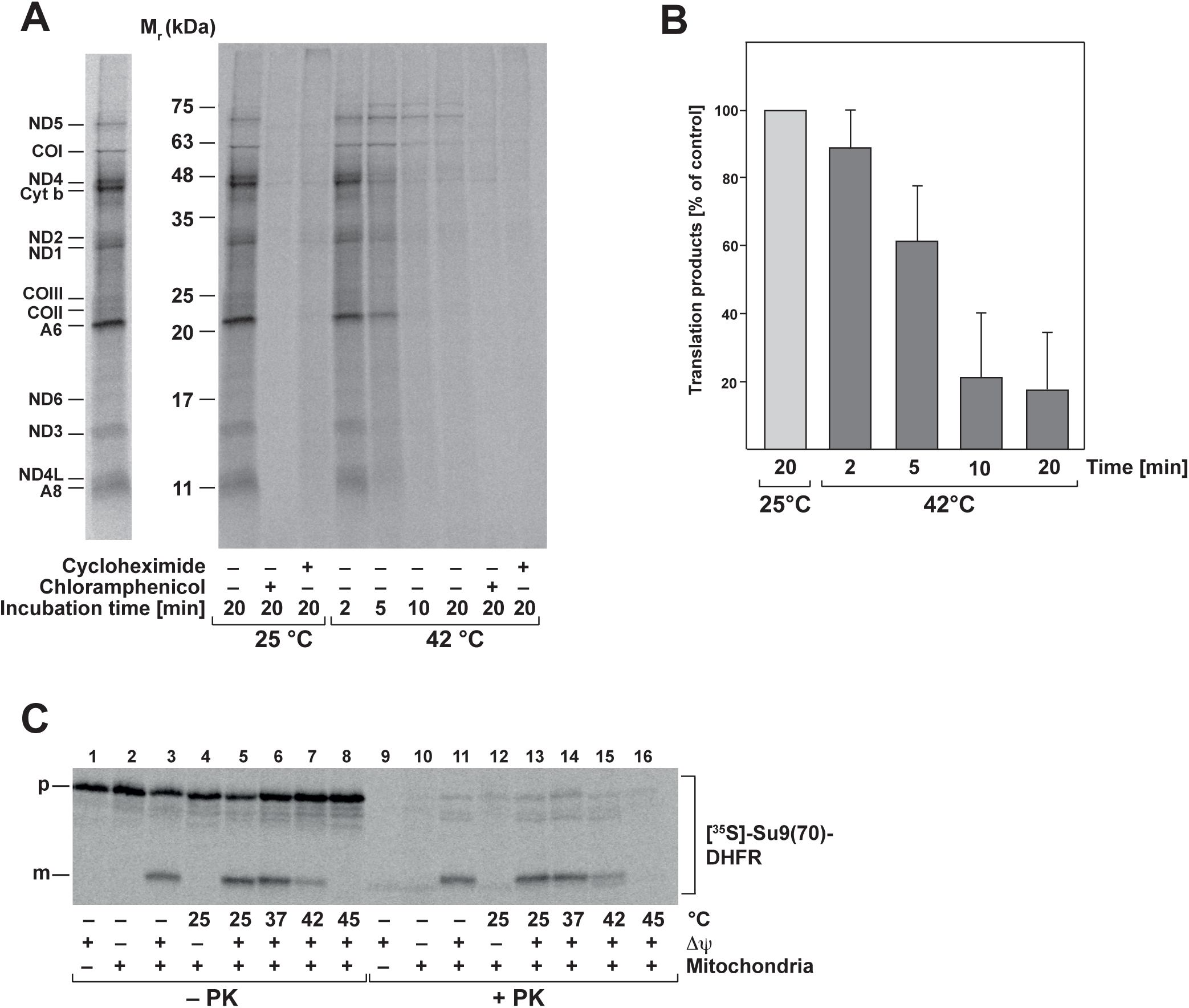
Mitochondrial protein biogenesis is compromised under heat stress. A) *In organello* translation of mitochondria-encoded proteins. Isolated mitochondria were heat stressed as indicated and newly translated proteins were labeled with [^35^S]-Met/Cys. Samples were treated with cycloheximide and chloramphenicol to impede cytosolic or mitochondrial translation. Left panel: Bands representing mitochondria-encoded subunits are indicated (ND, NADH dehydrogenase; CO, cytochrome *c* oxidase; A, ATP-synthase; Cyt b, cytochrome *bc1* complex). B) Quantitative analysis of *in organello* translation reactions as described in A). Shown are mean relative values and SEM of total incorporated radioactivity (n=3; control values at 25°C were set to 100%). C) Mitochondrial protein import after heat stress. *In vitro* import of [^35^S]-Met/Cys-labeled preprotein Su9(70)DHFR into isolated intact mitochondria after heat stress pre-incubation as indicated. Where indicated, the inner membrane potential (ΔΨ) was depleted. Non-imported preproteins were digested by incubation with proteinase K (PK). Translated or imported proteins were analyzed by SDS-PAGE and detected by digital autoradiography. p: precursor, m: mature protein.

As mitochondrial function depends both on endogenous translation as well as cytosolic translation of nuclear-encoded mitochondrial proteins and their subsequent import, we tested if there is also an impact of heat stress on the import efficiency in addition to the observed impairment of mitochondrial translation. We performed *in vitro* import experiments using an [^35^S]-labeled artificial mitochondrial preprotein Su9(70)-DHFR into fully energized isolated mitochondria at 30 °C after the mitochondria were subjected to a pre-incubation at heat stress temperatures for 20 min. A proteinase K treatment was used to remove non-imported preproteins. An import reaction with mitochondria where the membrane potential (ΔΨ) was depleted served as negative control. We observed that the import reaction was strongly influenced by heat stress (Fig. 6C). Whereas after the pre-incubation at 37°C the amount of imported protein was still comparable to the control condition, import efficiency decreased substantially after a heat shock at 42°C and was completely depleted after a severe heat shock at 45°C. This showed that after heat shock not only the new translation of mitochondrially encoded proteins broke down, but also no cytosolically translated proteins were imported into the mitochondria. As both processes are affected, this resulted in a complete shutdown of mitochondrial protein biogenesis under heat shock conditions.

We also determined if the newly synthesized mitochondrial proteins were prone to aggregate under heat stress. A translation reaction in intact isolated mitochondria was performed at 30°C as described above, generating radiolabeled mitochondrial nascent chains. After release of the nascent chains from the ribosome by treatment with puromycin, the mitochondria were subjected to a short heat-shock at 42°C. The formation of aggregates was tested by a detergent lysis, high-speed centrifugation, SDS-PAGE and autoradiography. Indeed, a high proportion of the nascent polypeptides were found in the aggregate pellet of mitochondria treated with 42°C (Fig. 7A). As control, we again observed a quantitative aggregation of Tufm and a partial aggregation of the mitochondrial Hsp90 chaperone Trap1, while membrane proteins like Tim23 were not found in the pellet fraction (Fig. 7B).

**Figure 7.**
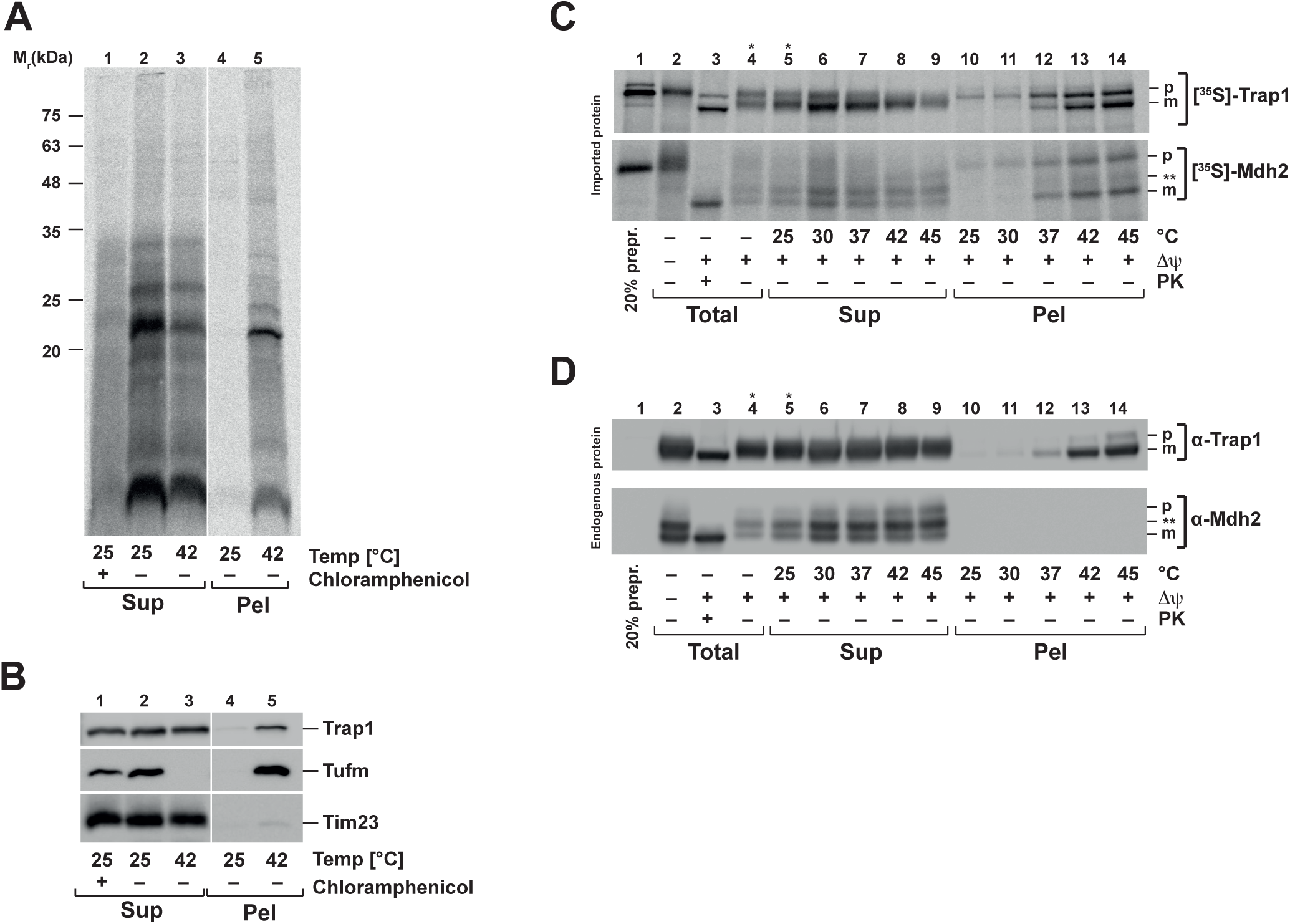
Heat-induced aggregation of nascent mitochondrial polypeptides. A) Heat stress of fresh synthesized mitochondria-encoded proteins. Newly translated mitochondrial proteins were labeled with [^35^S]-Met/Cys under normal conditions (30°C). Control samples were treated with chloramphenicol to impede mitochondrial translation. Directly after the translation reaction, mitochondria were heat-treated for 20 min at the indicated temperatures, lysed and aggregates were separated by centrifugation as described. Supernatants (Sup) and pellets (Pel) were analyzed by SDS-PAGE, Western blot and autoradiography. B) Samples prepared as described in A) were analyzed using the indicated antisera. C) Heat-induced aggregation of newly imported and endogenous proteins. After *in vitro* import of [^35^S]Met/Cys-labeled preproteins Trap1 and Mdh2 as described in Fig. 6C, mitochondria were heat-stressed for 20 min at indicated temperatures. After lysis, aggregates were separated by centrifugation and supernatants (Sup), pellets (Pel) as well as total amounts were analyzed by SDS-PAGE and Western blot (D) (ΔΨ, inner membrane potential; PK, proteinase K treatment). Translated or imported proteins were analyzed by SDS-PAGE and detected by digital autoradiography. p: precursor, m: mature protein. Asterisks indicate lanes where less sample was loaded.

We also performed an *in vitro* import of the genuine mitochondrial preproteins Trap1 (heat shock protein 75 kDa) and Mdh2 (malate dehydrogenase) into mitochondria to assess the aggregation propensity of newly imported preproteins. The import reaction was performed under normal temperature (30°C) as described. Directly after import, mitochondria were exposed to various temperatures to induce heat-stress. After lysis of mitochondria, aggregates were separated from soluble proteins by centrifugation and analyzed by SDS-PAGE. The newly imported radiolabeled preproteins Trap1 and Mdh2 were analyzed by autoradiography (Fig. 7C) while already present endogenous Trap1 and Mdh2 polypeptides were detected by Western blot (Fig. 7D). As the amount of the radioactive labeled imported proteins was too low to be detected by antibodies, both protein species could be distinguished. The freshly imported proteins were both aggregating in a temperature-dependent manner. After incubation at 37°C the imported Trap1 and Mdh2 partially accumulated in the pellet fraction although soluble polypeptides could still be detected in the supernatant. The amounts of aggregated preprotein increased strongly at 42°C and 45°C. The same behavior could also be detected for endogenous Trap1, which was also aggregating with rising temperatures (Fig. 7D). In contrast the endogenous Mdh2 showed no aggregation and stayed soluble even at incubation at 45°C. These observations indicated that both nascent proteins at the ribosome as well as newly imported proteins are highly sensitive to thermal aggregation independently of the thermodynamic stability of the native folding state under steady-state conditions.

## Discussion

As mitochondria represent semi-autonomous organelles within their eukaryotic host, their reactivity to diverse stress conditions has to be analyzed separately from the general cellular stress response. Therefore, we focused our study on the heat-sensitivity of mammalian mitochondrial polypeptides on a proteomic level. It has to be noted that heat stress had an obvious general effect on mitochondria as the overall mitochondrial shape in intact HeLa cells changed from the typical form of a tubular network, which is evenly distributed within the cell (24) to a more condensed structure concentrated near the nucleus. This observation confirms a previous electron-microscopic study in mammalian cells, which showed an increasing number of mitochondria at the nucleus in contrast to the cell periphery after heat stress, most likely due to a heat-dependent damage of the cytoskeleton (25).

In general, heat stress of mitochondria *in vivo* and *in vitro* caused major defects of typical biochemical properties, including a significant decrease of the inner membrane potential as well as a drop in mitochondrial ATP concentration. We also observed a small reduction in the amount of reactive oxygen species inside mitochondria, which most likely correlates with a decrease activity of the respiratory chain. This decrease in energetic efficiency occurred despite the overall composition and structure of the mitochondrial respiratory chain complexes as well as the integrity of mitochondrial membranes remained largely unaffected, demonstrating that mitochondria were not completely destroyed during the stress treatment. In addition, mitochondrial enzyme activities did not seem to be affected negatively by heat stress in general with the exception of the KGDHC. This key metabolic enzyme showed a small decrease in enzymatic activity, an observation that was previously made also in fungal mitochondria (26). Interestingly a decreased KGDHC activity was also observed in brain mitochondria of patients of Alzheimer disease (27), which could indicate an influence of aggregation of KGDHC in the onset of the disease.

The primary goal of this work was to characterize the impact of heat stress on the mitochondrial proteome in order to identify proteins, which are prone to misfolding and aggregation. Our analysis indicated a remarkable stress resistance of the mitochondrial proteome as most polypeptides stayed soluble after a treatment with elevated temperatures. Such an overall resistance against stress conditions was already revealed by a previous proteomic analysis of aggregating proteins in mitochondria from *Saccharomyces cervisiae* (15). In both studies just a small subset of mitochondrial proteins aggregated under heat stress conditions, supporting the idea that the heat sensitivity of a cell does not depend on a large-scale general denaturation and loss-of-function of proteins (28) but rather on the deactivation of few or single essential proteins with key physiological functions (29,30).

Although the overall protein amount in a soluble state was not affected strongly after temperature stress, a few mitochondrial proteins were also found in the aggregate pellet fraction. The most prominent example was the Krebs cycle enzyme aconitase, Aco2, previously identified as a meta-stabile polypeptide (15,31), Its heat sensitivity could be attributed to its complex structural composition, containing a Fe/S cluster as a large prosthetic group. A similar behavior was observed for the matrix chaperone Hsp60. As Hsp60 forms a homo-oligomeric protein complex consisting of 14 subunits, it may have an intrinsic sensitivity to different stress conditions. In addition, due to its chaperone properties it may also interact with other denatured polypeptides and hence be found in the aggregate pellet. Indeed, we observed that the AAA+ protease Lon homolog (LonP1), a protein exhibiting both chaperone and protease activities (32) was significantly reduced in solubility. The homolog of the protease LonP1, called Pim1, had been established as a major factor preventing a negative impact of heat-dependent aggregation by removing misfolded polypeptides (15,33). In this context, an association of at least a part of the Lon proteins with the misfolded polypeptides in the aggregate pellet could be expected.

In contrast, the mitochondrial translation elongation factor Tu (Tufm) was exceptionally sensitive to an exposure to elevated temperatures. Already under mild heat shock conditions Tufm was completely removed from the supernatant fraction by a genuine aggregation reaction. We also observed that the mitochondrial elongation factor Ts (Tsfm), a close partner protein and guanine nucleotide exchange factor for Tufm (20), showed the same heat-dependent aggregation behavior. The solubility of Tufm was also severely compromised under oxidative stress conditions, also representing an important stress factor for mitochondria (34). The significant thermo-lability of the mitochondrial elongation factors seems to be evolutionary conserved, as their bacterial relatives were originally labeled according to their thermostability as <u>u</u>nstable (u) and <u>s</u>table (s) (35,36). However more detailed later studies did not support this strong difference (37) and our results show that both elongations factors are highly thermolabile in mitochondria.

Interestingly, we did not observe a recovery of protein solubility when mitochondria were put back to normal temperature conditions, indicating that a chaperone-dependent disaggregation process is inefficient or even absent in mammalian mitochondria. Disaggregation has been demonstrated in fungal mitochondria under *in vivo* conditions (38,39), correlating with the presence of a specific member of the Hsp100 chaperone family, Hsp78, which is lacking in metazoan organisms. Although a principal disaggregation activity has been shown with mammalian chaperones in *in vitro* reactions (40,41), direct evidence for aggregate solubilization *in vivo* is still lacking for mammalian cellular systems.

Proteins from the elongation factor Tu family like Tufm are highly conserved proteins found in bacteria, plant chloroplasts, mitochondria and eukaryotic cytoplasm. Tufm, in cooperation with the nucleotid exchange factor Tsfm, facilitates the binding of aminoacyl-tRNA (aa-tRNA) to the acceptor site of the mitochondrial ribosome, thereby controlling translation progress in the elongation phase (42). Tufm, together with Tsfm, therefore represents a key element of the mitochondrial protein biosynthesis machinery. Hence, the exceptionally high thermo-lability of the translation factors Tufm and Tsfm and their complete deactivation by stress-induced aggregation suggested a major impact on the efficiency of the endogenous mitochondrial translation process at elevated temperatures. Indeed, the decreased solubility of the elongation factors of Tufm and Tsfm correlated very well with the decreased translation rate at elevated temperatures. This observation suggests that the thermo-labile elongation factors may serve as a stress-triggered switch to quickly shut-off mitochondrial synthesis of new polypeptides in order to prevent an accumulation of misfolded protein species. In *S. cerevisiae* mitochondria, the homolog of elongation factor Tu (Tuf1) was not identified as a prominently aggregating polypeptide (15). In these cellular systems, which contain an Hsp100-dependent dis–aggregation activity, a translational shutdown might not be required. In fact, the presence of an Hsp100 chaperone has been shown to enhance the recovery of fungal mitochondrial protein biosynthesis after heat stress (43).

In general, newly synthesized proteins exhibit an increased sensitivity to heat stress due to their incomplete folding state (44), an observation we also could confirm for mitochondrial nascent polypeptides. Stalling the biosynthesis of proteins during stress conditions would therefore be beneficial for the cell in terms of limiting the amount of potential aggregation-prone proteins. Indeed, a general heat stress-induced inhibition of protein synthesis was already described for cytosolically synthesized proteins in HeLa cells. The inhibitory effect is also exerted on the translational level and is caused by a phosphorylation of the translation initiation factor eIF2α (45,46). In addition, a heat-induced formation of stress granules in the cytosol, consisting of non-translated mRNA, RNA-binding proteins, translation initiation factors and 40S ribosomal subunits, has been described as a cellular mechanism for the prevention of an overloading of the protein quality control system by newly synthesized polypeptides (47). The formation of stress granules is also initiated by the block of translation initiation due to phosphorylation of eIF2α (48). An eIF2α-mediated translational shutdown has been also described as a major factor of the ER stress response (49).

A recovery from stress conditions includes a partial disassembly of stress granules as well as restoration of translation (47,50). However, in isolated mitochondria we were not able to detect a recovery of the solubility of aggregated translation factors Tufm and Tsfm as well as a recovery of the mitochondrial translation. However, our study focused on *in organello* conditions, where compensatory mechanisms provided by other cellular processes were excluded. The molecular details of potential recovery processes need to be addressed in future experiments.

Previous experiments indicated an involvement of the bacterial and plant homologs of the elongation factor (EF) Tu in heat stress reactions, although the postulated cellular mechanism underlying this effect was different. In addition to heat stress also other abiotic stresses like salinity, water restriction, and hypothermia resulted in upregulation of gene expression of both plastid EF-Tu as well as cytosolic EF-1α in different plant species (35,51), suggesting an important role in development of stress tolerance. A chaperone-like activity has been postulated for EF-Tu proteins from mammalian mitochondria (52), bacteria (53,54) as well as from plant chloroplasts. However, this hypothesis has been mainly based on experiments using the recombinant proteins in *in vitro* assays. Although there seems to be a correlation between a putative prevention of thermal protein aggregation by EF-Tu with improved heat tolerance in plants (55,56), so far it is not clear, how relevant a putative chaperone-like activity is in the *in vivo* situation.

An important aspect of mitochondrial protein biogenesis is the fact that only a minority of the resident proteins is provided via mitochondrial translation. The vast majority of mitochondrial proteins is encoded in the nucleus and imported after their synthesis in the cytosol (57). Hence, an unilateral shut-down of mitochondrial translation would only exacerbate the problem of an accumulation of misfolded polypeptides inside mitochondria. Interestingly, we observed under *in vivo* like conditions that the import efficiency of cytosolic proteins was strongly decreased at elevated temperatures. This correlated with the observation that similar to nascent mitochondrial proteins at the ribosome, also newly imported polypeptides are highly sensitive to heat stress and prone to aggregation. Although a mitochondrial unfolded protein response (mtUPR) has been described as a potential protective mechanism against the accumulation of misfolded mitochondrial polypeptides (58), no specific effect of this stress-regulatory process on mitochondrial protein biosynthesis has been described so far. As we did not observe induction of stress-dependent mitophagy under the used conditions (Fig. S6), prevention of potential proteotoxicity seems to depend largely on internal mitochondrial mechanisms. Thus, a translation shutdown by Tufm aggregation in combination with a decrease in import efficiency would effectively prevent an accumulation of misfolded polypeptides in mitochondria under heat stress – representing a protective mechanism to prevent further proteotoxic insults on organellar function. Hence, our observations support the notion that organellar EF-Tu proteins represent important components of the stress-protective cellular processes responsible for protein homeostasis.

## Experimental Procedures

### Cells and culture conditions

HeLa cells were cultured in Roswell Park Memorial Institute medium (RPMI medium), supplemented with 10% (v/v) fetal calf serum (FCS), 2 mM L-glutamine, 100 units/ml penicillin, 100 μg/ml streptomycin. Cells were grown in tissue culture dishes at 37°C in 5% CO2 and routinely passaged by trypsinization at ratios 1:3 to 1:6 every 48 to 72 h.

### Flow cytometry

Membrane potential was assayed by an incubation of cultures cells for 10 min at 37°C with 0.5 μM tetramethylrhodamine ethyl ester (TMRE). As control, a sample was treated with valinomycin (f. c. 0.1 μM) for 30 min at 37°C. The generation of ROS was determined using the mitochondrial superoxide indicator MitoSOXTM (Thermo Fisher Scientific). Cells were incubated with the indicator dye for 10 min at 37°C. As positive control, cells were treated with 100 μM menadione for 15 min at 37°C. Cells were harvested by trypsinization and washed twice in PBS containing 0.2% (w/v) BSA. Fluorescence of 20,000 cells for each sample was analyzed using the flow cytometer CyFlow space CY-S3001 (Partec).

### Mitochondrial aggregation assay

Mitochondria were isolated from cultures cells essentially as described (59). For each sample 30 μg of fresh mitochondria were resuspended in 100 μl resuspension buffer (500 mM sucrose, 160 mM potassium acetate (KAc), 40 mM HEPES/KOH pH 7.6, 10 mM MgAc, 5 mM glutamate, 5 mM malate, 1 mM DTT). For analysis of the nucleotide dependence the resuspension buffer was supplemented with 5 mM adenosine triphosphate (ATP), creatine phosphate (20 mM) and creatine kinase (4 μg/ml). Analysis of the GTP-dependence was performed in detergent-lysed mitochondria (lysis buffer: 0.5% (v/v) Triton X-100, 200 mM KCl, 30 mM Tris/HCl pH 7.4, 0.5 mM phenylmethyl– sulfonylflouride (PMSF), 1x protease inhibitors) supplemented with 10 mM MgAc and 2.5 mM GTP. Mitochondria were either pre-incubated for 20 min at indicated temperatures (25°C-45°C) or in presence of menadione (0-2 mM) at 25°C. After incubation, mitochondria were reisolated by centrifugation (12,000 x*g*, 10 min, 4°C) and pellets were lysed in lysis buffer containing 5 mM EDTA, by vigorous shaking for 10 min at 4°C. Soluble proteins were separated from aggregated proteins by centrifugation at 20,000 x*g* for 20 min at 4°C. The supernatants were precipitated with TCA and pellets were re-extracted in 100 μl lysis buffer by shaking as described above. After a second centrifugation step at 20,000 x*g* for 20 min at 4°C supernatants were discarded and the pellet was resuspended in 1x Laemmli SDS-PAGE buffer (2% (w/v) SDS, 10% (v/v) glycerol, 60 mM Tris/HCl pH 6.8, 5% (w/v) β-mercaptoethanol, 0.02% (w/v) bromophenol blue). Samples were analyzed by SDS-PAGE and Western blot.

To distinguish between aggregate sizes, different centrifugation steps were applied. After the first low spin step at 1,600 x*g* for 5 min the supernatant was collected and centrifuged at 20,000 x*g* for 5 min, following a 20 min centrifugation at the same spin. After the last spin at 125,000 x*g* for 40 min the supernatant was precipitated with TCA. All pellets were resuspended in 1x Laemmli buffer and analyzed as described above.

For recovery from aggregation, 30 μg of mitochondria were resuspended in 100 μl incubation buffer (250 mM sucrose, 80 mM potassium acetate, 20 mM HEPES/KOH pH 7.6, 5 mM magnesium acetate, 5 mM glutamate, 5 mM malate, 5 mM KPi pH 7.4, 2 mM ATP, 1 mM DTT) to keep them fully energized. After heat stress for 20 min at 42°C samples were incubated for up to 2 h at 30°C. Samples were lysed, centrifuged and analyzed as described above.

### Proteinase K treatment of mitochondria

For determination of the integrity of mitochondria, mitochondria were treated after heat stress with 1.25 μg/ml proteinase K for 20 min. Treatments were stopped by 15 min incubation with 1.5 mM PMSF and following precipitation by TCA. To assess proteinase resistance of polypeptides, mitochondria were lysed with lysis buffer containing 5 mM EDTA and incubated with proteinase K. Protease treatment was stopped by adding PMSF and samples were precipitated with TCA.

### Membrane potential measurement

30 μg mitochondria were resuspended in potential buffer (600 mM sorbitol, 20 mM KPi, pH 7.2, 10 mM MgCl2, 10 mM glutamate, 5 mM malate, 0.1% (w/v) BSA) and incubated with 1 μM TMRE for 10 min at 30°C. Negative control samples were treated with valinomycin for 10 min at 30°C prior TMRE incubation. All samples were washed with potential buffer and TMRE fluorescence (excitation: 540 nm, emission: 585 nm) was measured in a microplate reader (Infinite M200 PRO, Tecan).

### Determination of ATP content in isolated mitochondria after heat shock

The mitochondrial ATP content was measured by a luciferase based ATP assay (ATP Determination kit, Invitrogen). Isolated energized mitochondria were either analyzed directly or after 20 min incubation at 25°C or 42°C. The ATP assay was performed according to the manufacture’s instructions and luminescence was measured in a microplate reader.

### Enzymatic assays

Activities of malate dehydrogenase (MDH) and alpha-ketoglutarate dehydrogenase complex (KGDHC) were measured in isolated mitochondria (100 μg/sample for MDH and 200 μg/sample for KGDHC activity assay) after incubation for 20 min at 25°C (control) or 42°C (heat shock) in resuspension buffer. After reisolation of the mitochondria at 12,000 x*g* for 10 min at 4°C, the mitochondrial pellets were lysed in 50 μl homogenization buffer (0.5% Triton X-100, 50 mM Tris/HCl pH 7.4) and shaken at maximum speed for 10 min at room temperature. The respective reaction mixtures (MDH activity assay: 50 mM Tris/HCl pH 7.4, 0.5 mM oxalacetic acid and 75 μM NADH; KGDHC activity assay: 50 mM Tris/HCl pH 7.4, 50 mM KCl, 5 mM alpha-Ketoglutarate, 2 mM MgCl2, 0.3 mM thiamine pyrophosphate, 0.3 mM coenzyme A and 2 mM NAD+) were pre-warmed at 37°C, the lysed mitochondria were added and enzyme activities were measured at 340 nm every 15 sec for 10 min in a photometer (BioPhotometer plus, Eppendorf) by either decrease of NADH absorption (MDH) or increase of NADH absorption (KGDHC).

### Blue Native SDS-PAGE

40 μg of isolated mitochondria were resuspended in resuspension buffer, incubated for 20 min at 25°C (unstressed) or 42°C (heat stress) and afterwards solubilized in digitonin lysis buffer (0.4% (w/v) digitonin, 50 mM NaCl, 20 mM Tris/HCl, pH 7.4, 2 mM EDTA pH 8.0, 1 mM PMSF, 10% (v/v) glycerol) by pipetting (20x) and shaking for 10 min at 4°C at maximum speed. 10x loading dye (5% (w/v) Coomassie blue G–250, 500 mM ε-amino n-caproic acid, 100 mM Bis-Tris-HCl pH 7.0) was added and samples were applied to a 5-16% polyacrylamide gradient gel. The samples were analyzed by Western blot using antibodies against subunits of the respiratory chain complexes as indicated.

### 2-D gel electrophoresis

Isolated mitochondria (150 μg total protein) were dissolved in resuspension buffer and incubated for 20 min at 25°C (control) or 42°C (heat stress). After centrifugation at 12,000 x*g* for 10 min at 4°C mitochondria were lysed in lysis buffer (0.5% Triton X-100, 200 mM KCl, 30 mM Tris/HCl pH 7.4, 5 mM EDTA, 0.5 mM PMSF, 1x protease inhibitors) by vigorous shaking for 10 min at 4°C. The aggregated proteins were separated ultracentrifugation at 125,000 x*g* for 40 min. Supernatants were precipitated with TCA and re-suspended in IEF lysis buffer (7 M urea, 2 M thiourea, 4% (w/v) CHAPS, 10 mM DTT, 1x protease inhibitors, 40 mM Tris/HCL pH 8.8) and lysed by 10 min of shaking at maximum speed at 4°C. Control, heat stressed and standard samples were labeled with CyDye DIGE Fluor minimal dyes (GE Healthcare). The pooled samples were diluted in rehydration buffer (7 M urea, 2 M thiourea, 2% (w/v) CHAPS, 20 mM DTT, 0.8% (v/v) PharmalyteTM 3–11 NL, small amount of bromophenol blue) and applied to a 18 cm non-linear, pH 3–11 ImmobilineTM Dry Strip (GE Healthcare) in an IPGphor isoelectric focusing system (GE Healthcare). In parallel, one sample containing 400 μg total mitochondria was precipitated with TCA, resuspended in rehydration buffer and applied to a strip as described above. Protein spots were detected by Coomassie blue G–250 staining and manually excised for analysis by mass spectrometry.

Immobiline strips were rehydrated ingel rehydration for 12 h at 30 V. Afterwards protein focusing was conducted stepwise for respectively 1 h at 200 V, 500 V and 1000 V followed by a slow increase (2 h and 30 min) to 8000 V for 3 h and finally decrease to 500 V for 2 h and 30 min. Focused strips were incubated in equilibration buffer (6 M urea, 1.5 mM Tris/HCl pH 8.8, 50% (v/v) glycerol, 1% (w/v) SDS) containing 1% (w/v) DTT for equilibration followed by an incubation in equilibration buffer containing 2.5% (w/v) iodacetamide for reduction. Both incubations were performed for 15 min in the dark. Proteins were separated by SDS-PAGE on a 11% polyacrylamide gel for 1 h at 0.5 W per gel followed by 1.5 W per gel over night at 20°C in an Ettan^TM^ DALT*six* system (GE Healthcare). Proteins were detected by fluorescence scanning using the Fluoro Image Analyzer FLA-5100 (Fujifilm) and obtained protein spots were analyzed by the Delta2D software (DECODON). For each gel spot intensities were normalized by the total sum of spot intensity at 42°C divided by the total sum of spot intensity at 25°C and the average of six different gels as well as the standard error of the mean (SEM) was calculated. The significance was calculated using GraphPad Prism 6 software via 1-way ANOVA of all samples to a control of 100% (Dunnett’s multiple comparisions test). A p–value ≤ 0.001 was considered significant and indicated by asterisks. Protein spots of interest were manually excised and analyzed by mass spectroscopy (see supporting experimental procedures).

### Preprotein import into mitochondria

*In vitro* import was performed as described (59), using the radiolabeled substrate proteins Su9(70)-DHFR, Trap1 and Mdh2, which were synthesized by *in vitro* transcription and translation in rabbit reticulocyte lysate. For each import reaction 50 μg of fresh isolated energized mitochondria were used that were either heat-shocked before the import reaction to assess tha import efficiency or after the import reaction to test if fresh imported proteins were prone to aggregate. The heat shock was performed in resuspension buffer for 20 min at respective temperatures. Control samples without membrane potential (ΔΨ) mitochondria were pretreated with antimycin, valinomycin and oligomycin. The import reaction was performed for 40 min at 30°C and stopped by adding 0.5 μM valinomycin. Where indicated, samples were treated with 50 μg/ml proteinase K for 30 min at 4°C to digest non-imported proteins. Samples were analyzed by SDS-PAGE and autoradiography.

### *In organello* translation

50 μg of fresh isolated mitochondria were resuspended in resuspension buffer and incubated 2-20 min at 42°C and 25°C. After reisolation of the mitochondria (12,000 x*g*, 10 min, 4°C) pellets were resuspended in translation buffer (645 mM sorbitol, 160 mM KCl, 21.5 mM Tris/HCl pH 7.4, 21.5 mM ADP, 13.5 mM MgSO4, 3.2 mg/ml BSA, 14 μM amino acid mix w/o methionine, 0.53 mM GTP, 180 mM creatine phosphate, 4 μg/ml creatine kinase, 1.2 mg/ml α-ketoglutarate). Where indicated cycloheximide (f. c. 100 μg/μl) or chloramphenicol (f. c. 100 μg/μl) were added. After pre–incubation of the samples for 3 min at 30°C and shaking at 300 rpm, newly synthesized mitochondrial proteins were labeled by adding [^35^S]-methionine/ cysteine (f. c. 22 μCi/μl). The samples were incubated for 45 min at 30°C and labeling was stopped by adding MOPS/Met buffer (1 M MOPS, pH 7.2, 200 mM methionine, f. c. 50 mM) and incubating for 30 min at 30°C. Mitochondria were washed once with washing buffer (0.6 M sorbitol, 1 mM EDTA, 5 mM methionine). For further analysis mitochondria were lysed in lysis buffer (see above) containing 50 μg/ml (f. c.) puromycin and aggregates were isolated after heat stress as described before. Samples were analyzed by SDS-PAGE on a 15% polyacrylamide gel containing 1.1 M urea and autoradiography.

## Acknowledgements

We are grateful to M. Fuhrmann and B. Gehrig for expert technical assistance. We thank W. Jaworek, K. Pollecker and Dr. G. Cenini for discussion and critical comments on the manuscript. Work in the authors laboratory was supported by the Deutsche Forschungsgemeinschaft (grant VO 657/5-2 to W.V.).

